# Optimizing strain selection for association studies under hard cost constraints

**DOI:** 10.1101/2025.05.31.657208

**Authors:** Christoph D. Rau, Patrick H. Bradley

## Abstract

Quantitative genetics methods that link genotype to phenotype can be especially powerful tools in model organisms and non-human populations, where researchers can apply well-controlled perturbations to commercially-available collections of strains. However, purchasing and phenotyping large collections can be cost-prohibitive. We evaluate several approaches to select subsets of a panel of strains with the aim of maximizing power under fixed budgetary constraints. Some approaches focus solely on costs, others on genetic diversity, and some on both simultaneously. To assess how these results generalize, we simulate data across two biological and statistical settings: phylogenetic regression across bacterial isolates, and linear-mixed model regression across mouse strains. Surprisingly, we find that simply selecting the cheapest strains until the budget is exhausted (MinCost) is usually among the top-performing methods, and that algorithms that consider genetic diversity without weighting by cost tend to show much lower power overall. Methods that consider both cost and diversity are typically, though not universally, the top performers. A comparison of diversity-maximizing objectives reveals that the most commonly studied objective (MaxMin) tends to be the among the least effective at retaining power. Instead, two alternative objectives that have not been previously studied in this context — maximizing the total minor allele frequency (MaxMAF) and maximizing the total sum of pairwise genetic distances (MaxSum) — typically perform best. Finally, in addition to fully simulated data, we also test these approaches by subsampling real data on cardiovascular phenotypes from the Hybrid Mouse Diversity Panel (HMDP). The three best methods on these data are cost-aware (p-Median, MaxSum, and MinCost). Based on these results, we recommend investigators consider approaches that maximize total MAF or pairwise genetic distance weighted by strain cost, but we also conclude that a surprisingly robust approach is to simply pick the most strains one can afford.

## INTRODUCTION

The cost of research has grown significantly over the past two decades. One measure is the average size of an R01 grant from the National Institutes of Health, which has increased 87% between 1999 and 2019^1^, 64% faster than inflation. These cost increases come from many factors: novel techniques and technologies that supplant other, often cheaper approaches (e.g. PCR to RNA microarrays to RNAseq to single cell RNAseq), more complex study designs which necessitate additional research cohorts and endpoints, and, notably, a drive to increase the size of studies to attempt to both achieve greater statistical power and identify less penetrant drivers of disease conditions^2–4^. For example, over the past two decades, the GWAS community has been gathering larger and larger populations to try and recover the effects of polymorphisms with smaller and smaller effect sizes^5,6^. Depending on how they are funded, such large populations may be out of reach for individual investigators. Furthermore, beyond simple financial considerations, researchers who use animal populations are often mandated to follow the three Rs (Replacement, Reduction, Refinement) in their research, which states that if it is possible to achieve the same or similar result with fewer animals, then researchers should make every effort to do so; for example, the US Food and Drug administration has recently announced a roadmap to reduce animal testing in preclinical studies^7^. Taken together, the increasing costs in both money and time of doing research coupled with ethical responsibilities to animal cohorts are powerful incentives to optimize and streamline research projects, as long as this does not come at the expense of statistical power.

In comparative studies, one opportunity for streamlining concerns how we select the strains that will be phenotyped. Diverse collections of microbes have proven to be powerful tools for finding genetic variants associated with phenotypes including host range of pathogens^8,9^, positive fitness effects of symbionts^10^, toxin tolerance in industrially-relevant microbes^11^, and gene and protein expression in model organisms^12^. When applied in diverse animal lines, GWAS, MPRA assays, and other comparative omics-scale analyses have allowed us to link genetic polymorphisms to complex disease susceptibility^13–18^. However, prices for such strains are not uniform: rather, they vary widely across strain collections, individual genotypes, and even how a particular strain is propagated and shipped. For example, a mouse from Jackson Labs costs a different amount of money depending on its strain, age, and sex, ranging from less than $30 for a 3-week-old C57BL/6J male, to over $260 for a 9 week old NZO/HILtJ male, to nearly $2,500 a litter for one of the many cryopreserved strains available for purchase. Similar factors apply to bacterial strains. Type strains are required to be preserved in at least two repositories, which may have different costs: the type strain of *Lachnospira eligens*, shipped as lyophilized cells, costs $474 from the American Type Culture Collection (ATCC) but 100 EUR (∼$113) from the German Collection of Microorganisms and Cell Cultures (DSMZ). As with mice, the format also matters: a live, active culture of the *L. eligens* type strain from DSMZ would cost 240 EUR ($273). Finally, both ATCC and DSMZ, as well as many other repositories, have unique strains that can only be purchased from them. Being able to optimize the selection of strains to maximize statistical power for a given budget would be a major benefit for the labs which use these populations.

One well-researched framing of strain selection, which Weitzman named the “Noah’s Ark” problem^19^, comes from bioconservation. Solutions to this problem maximize a measure of community diversity, typically either Faith’s phylogenetic diversity or PD (here, we refer to these as MaxPD methods)^20^, or the minimum pairwise distance between all selected taxa (which we term MaxMin)^19^. In the equal costs setting, MaxMin is equivalent to the p-dispersion problem from operations research, first formulated in the context of locating facilities on a network^21^. The general p-dispersion problem is NP-hard, but greedy MaxMin algorithms to select taxa perform well^22^, and under certain assumptions (distances given by an ultrametric tree), they also maximize the PD of the selected taxa. In fact, greedy MaxPD algorithms, where successive strains are chosen iteratively based on which additional strain leads to the largest increase, provably reach the global optimum^23,24^. Greedy methods to maximize PD have also been applied in genotype imputation studies, where they have been shown to improve power relative to random selection^25,26^.

However, while Weitzman’s original “Noah’s Ark” problem does consider unequal costs, most studies on strain selection in the context of association or genomic studies do not, and it is less clear how well greedy methods perform in this setting. Weitzman provides one way to integrate cost information: a greedy search where strains are ranked by genetic distinctiveness divided by cost^27^, which he terms the “myopic rule.” A similar approach, proposed by Hartmann and Steel^19^, is to maximize the sum of the PD and a penalty term, which in turn is a sum of some function of the costs of the selected strains. In this formulation, the greedy algorithm is still guaranteed to be optimal^23^ in the sense that it does maximize this modified objective. However, this may not be the same as directly optimizing PD under fixed costs. In the extremes, a function that multiplies all costs by zero would be equivalent to not considering costs at all, while multiplying all the costs by a very large number (relative to the maximum possible PD) would be equivalent to prioritizing the cheapest strains without considering diversity. Pardi and Goldman^28^ did develop an efficient, pseudo-polynomial time algorithm for directly maximizing PD under cost constraints, meaning that out of all sets of taxa that can be purchased with a given budget, the set with the highest PD will be chosen. However, this algorithm involves decomposing the problem by clade, so the taxa being considered must be related by a phylogenetic tree. Maximizing diversity on a tree via PD has many advantages, both in terms of constraining a more general NP-hard problem into one that can be solved exactly, and also in terms of achieving bioconservation outcomes^29,30^. However, metrics like PD cannot capture genetic relationships between taxa that are not described well by a tree: for example, when there is extensive horizontal transfer between species, as with gut bacteria^31^, or when there is a complex kinship matrix, as in panels that derive from backcrosses. Because these relationships may be particularly relevant in the context of genome-wide association studies and comparative genomics, the current study focuses on methods that can use more general measures of genetic distance between strains.

Furthermore, even assuming we could successfully maximize genetic divergence, doing so may not necessarily accomplish the aim of conserving statistical power^32^. For example, McAuliffe and colleagues ^32^developed an approach for comparative genomics in which they directly maximized power. They showed that the selected taxa were often very different from those that maximized evolutionary divergence, having as few as one taxon in common. Choosing power itself as the quantity to maximize has obvious advantages; however, this method requires calculating power in a combinatorial search over possible subtrees. While McAuliffe et al. derived an exact analytical solution for the power of their test, making it much more tractable, the computational cost of applying this method in a panel of hundreds or thousands of strains would be prohibitive.

Other metrics besides PD or MaxMin diversity could potentially offer better performance. For example, Matsen IV et al^33^ developed an approach that <u>min</u>imizes the <u>a</u>verage phylogenetic <u>d</u>istance of query sequences to the <u>c</u>losest leaf on a reference tree (“minimum-ADCL”), essentially maximizing how “representative” the selected subset is of the whole panel. As the authors note, this is a variation of the k-medoids problem^34^ on a tree, and as elaborated by Faith, it is also equivalent to discrete p-median optimization of diversity^35^. The p-median problem (closely related to k-medoids) can be framed as an integer linear programming problem. Kang et al. adapted Matsen’s p-median approach to pick reference panels for genotype imputation, and showed that for this particular problem, p-median outperformed both random and maximizing total diversity approaches^36^. However, these p-median methods do not consider costs and would have to be modified for the question we are interested in. Additionally, a p-median approach should deprioritize genetic outliers^33^; in the setting of quantitative genetics, however, outliers might actually improve the power to detect associations with rare variants.

Beyond MaxPD, MaxMin and p-median, there are also other approaches that could be used to maximize diversity^37^ that have not been explored to the same extent in the genetics literature. One of these is maximizing the sum of the total genetic distances between all selected strains, known as the MaxSum or (discrete) p-dispersion-sum problem^21,38^. Solutions to this problem can differ markedly from p-dispersion or p-median solutions (for instance, MaxSum solutions may include pairs of points that are close together^21^), potentially translating into different effects on power. Again, however, this problem is NP-hard, and most available heuristics have been developed for a formulation that does not include unequal costs.

Finally, because genome-wide association studies typically show the most power for common variants, instead of maximizing distances, one might also consider choosing the strains that maximize the minor allele frequency (or MAF, which ranges from 0 to 0.5) of all alleles in a panel, which by analogy we call MaxMAF. A related approach from crop science, called the “M strategy”^39,40^, aims to choose strains that maximize total allele diversity across a set of genetic markers; compared to methods based on genetic divergence, this strategy selects taxa bearing fewer non-informative alleles^41^. To our knowledge, however, neither MaxSum nor MaxMAF has been assessed in terms of retaining power in gene association studies, and these methods also have yet to be adapted to the setting of unequal costs.

In this manuscript, we set out to study different approaches for strain selection under a hard cost constraint, and to quantify how these methods affect our power to detect genetic associations (Figure 1). Because we wanted to apply these methods in settings where a tree might not best describe the genetic relationships between strains, we did not further consider methods for maximizing PD, and instead focused on approaches that allowed more flexible measures of genetic distance. Here, we consider five families of methods for considering genetic diversity:

- <u>None</u>: Only the number of strains is considered;
- <u>p-Median</u>: Minimize the total minimum genetic distance between non-selected and selected strains (as with the min-ADCL method);
- <u>MaxMin</u>: Maximize the minimum genetic distance between selected strains;
- <u>MaxSum</u>: Maximize the total genetic distance between all pairs of selected strains;
- <u>MaxMAF</u>: Maximize the sum of minor allele frequencies (MAFs).

**Figure 1:**
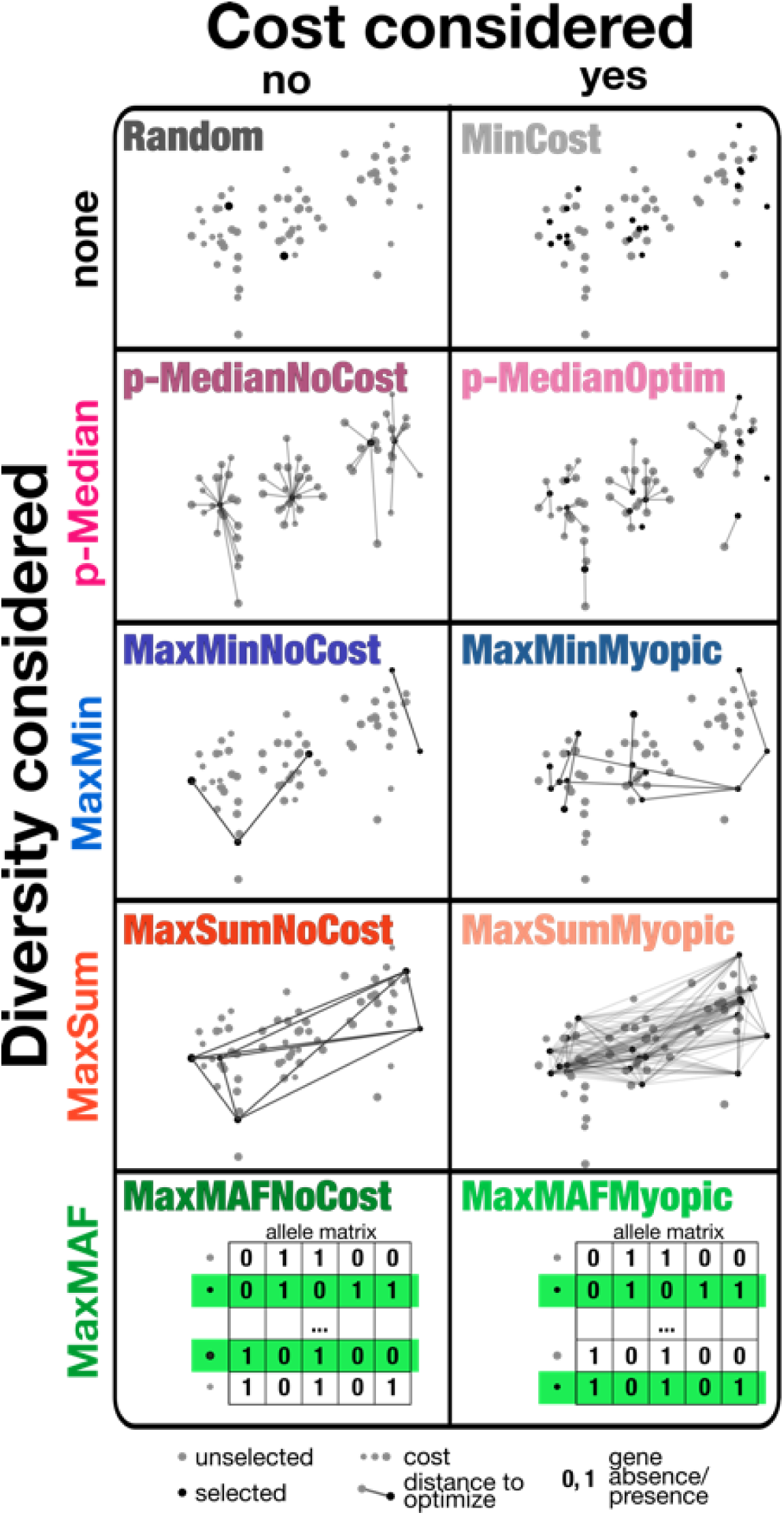
Illustration of methods considered for strain picking. Each method is illustrated with an example where selected strains are represented by black circles and unselected strains by gray circles, with cost being represented by the circle size. Methods in the same column either do not (left) or do (right) consider the cost of strains, while methods in the same row either do not consider diversity (top, Random and MinCost) or consider it in the same way (other rows). The p-Median, MaxMin, and MaxSum methods all optimize a function of distances between strains (gray lines connecting circles). The p-Median methods (purple) minimize the total distance between unselected strains and the nearest selected strain. The MaxMin methods (blue) maximize the minimum distance between selected strains, while the MaxSum methods (red) instead maximize the total distance. Instead of distances, the MaxMAF methods maximize the sum of minor allele frequencies, shown by the inset matrices. “Myopic” indicates a greedy search that prioritizes strains by diversity divided by cost, while “Optim” indicates a method that directly optimizes the objective using integer linear programming (ILP). The “NoCost” methods also use greedy search, except for “p-medianNoCost” which runs the Partitioning Around k-Medoids algorithm with successively greater *k*.

For each family, we test an algorithm that does not consider cost (“NoCost”), simply stopping when the budget is exhausted, and at least one algorithm that does (denoted by either “Myopic” or “Optim”). The “NoCost” algorithm is a greedy search, taking the strain that improves the family objective the most at each step, except in the case of “p-MedianNoCost,” where greedy search is known to perform poorly and we instead use the Partitioning Around Medoids algorithm. The “Myopic” algorithms we consider are greedy searches that divide a diversity measure by the cost of the strain, as with the “myopic rule.” Finally, the “Optim” algorithm directly optimizes the objective via mixed-integer linear programming, or MILP. (Note that while we did implement versions of MaxMinOptim and MaxSumOptim, the only MILP method that was computationally tractable on our data proved to be the p-MedianOptim algorithm.)

To our knowledge, this is the first systematic test of how different methods for cost-constrained strain selection affect power in downstream gene association studies. Additionally, the few studies to consider costs have focused on the MaxPD and MaxMin objectives; we have not found any studies that have implemented and tested cost-aware versions of the alternative MaxSum, MaxMAF, or p-Median approaches.

We evaluate these methods using simulations, which we conduct in two different settings: phylogenetic regression in microbes, and GWAS in mice. In the first setting, we model the effects of strain selection on power in phylogenetic regression applied to synthetic data from genotyped bacterial isolates. We perform two tests, one where we use the real genotypes of isolates from *Lachnospiraceae*, the most prevalent gut bacterial family worldwide, and one where we introduce a form of study bias by simulating an additional oversampled clade. In the second setting, we model the effects of strain selection on GWAS results in the Hybrid Mouse Diversity Panel, a commonly used genetic reference population for studying a number of phenotypes and diseases^42–46^. In our GWAS simulations, we vary the minor allele frequency (MAF) and effect size cutoffs (χ^2^), and also explore the impact of reducing the variance of prices by adding extra equal, fixed costs per strain (as might be appropriate when the cost of strain maintenance and/or phenotyping is a non-negligible portion of a research plan’s budgetary constraints). Finally, we follow up our simulation studies with an analysis of actual data from an HMDP GWAS study in which the beta adrenergic agonist isoproterenol was used to induce cardiac hypertrophy, which we subsample to simulate subset selection.

## METHODS

### Data Sources

Mouse genotype data from the Mouse Diversity Array for the Hybrid Mouse Diversity Panel (HMDP) was obtained from the Mouse Genomics Informatics website^17^. Mouse phenotype data is from a previously published dataset^18^ and is available at Mendeley Resources at data.mendeley.com/datasets/y8tdm4s7nh/1^47^. Mouse costs were pulled from the Jackson Labs website^48^ as on 4/05/2024 and represent the cost of 1 male and 2 female mice or one litter of cryopreserved animals, depending on the current status of the animals at the time of accession.

Metadata for bacterial strains was downloaded from BacDive^49^ as of 07/31/2025. Strains with a genome in NCBI’s GenBank or RefSeq were retained, then matched with the metadata from the Genome Taxonomy Database (GTDB)^50^ release 220 (downloaded 10/31/2024) to obtain taxonomic assignments, ignoring genome version numbers. Genomes were then filtered for GTDB annotations for the family Lachnospiraceae. The five most common strain repositories for these genomes were the German Collection of Microorganisms and Cell Cultures (DSMZ), Japan Collection of Microorganisms (JCM), American Type Culture Collection (ATCC), Culture Collection University of Gothenburg (CCUG), and Korean Collection for Type Cultures (KCTC). Costs for each repository were estimated by spot-checking five of the *Lachnospiraceae* strains on the respective websites and averaging the results; shipping costs were not included. When the exact same strain was available in multiple collections, we retained the cheapest copy. We used OrthoFinder^51^ to identify orthogroups (i.e., genes descending from a common ancestor in this group of species). Orthogroups with identical distributions across genomes were merged into phylogroups, then filtered such that all remaining phylogroups were present in at least three genomes and absent in at least three genomes.

### Simulation Studies

#### Phylogenetic regression

We performed two sets of tests: one using real *Lachnospiraceae* genotypes and the real tree (“Real”), and one using simulated genotype data on a version of the real *Lachnospiraceae* tree that had been augmented by a set of 100 very similar strains with very low cost (“Augmented”), to mimic a situation where one taxon may be highly oversampled relative to all others.

#### Real

We started with the *Lachnospiraceae* genome tree generated by OrthoFinder, making it ultrametric by using relative evolutionary divergence normalization^52^ using the “castor” package. For each scenario, we simulated 5,000 phenotypes on this ultrametric genome tree under a Brownian motion model using the “rTrait” function in the “phylolm” package. For each phenotype, we picked a phylogroup (without replacement), and then added an offset to that phenotype when the phylogroup was present, with the offset corresponding to 60% of that phenotype’s standard deviation. Phylogroups present in fewer than three genomes or absent in fewer than three genomes were excluded. We finally performed one phylogenetic regression per phenotype and phylogroup pair, allowing for measurement error, using the phylolm function in the “phylolm” package. We estimated power as the proportion of these tests with a p-value less than 0.05, replacing any tests that returned an NA value (as might happen when, e.g., there is no variance in the phylogroup) with a p-value of 1.

#### Augmented

Starting with the above tree of *Lachnospiraceae* genomes, we first grafted a random coalescent tree of 100 strains onto the immediate ancestor of *Agathobacter rectalis*, with the random tree scaled to 1/10^th^ the node depth of this ancestor. The presence/absence vectors of 5,000 synthetic phylogroups were then simulated using the rBinTrait function in phylolm^53^, which uses a continuous time Markov chain approach to model binary traits along a tree. For each synthetic phylogroup, the parameters alpha and beta were obtained by performing Ives-Garland phylogenetic logistic regression^54^ on one of the 5,000 phylogroups selected in the “real” simulation, using the phyloglm function in the phylolm package^54^. We used rejection sampling to make sure that all simulated phylogroups were present in at least three genomes and absent in at least three genomes. Finally, phenotypes were simulated and power was assessed as in the “real” simulation.

#### Recombinant

To assess the impact of genetic structure on the results, we first selected a set of six “parental” *Lachnospiraceae* strains using PAM^55^, then added a set of 100 simulated strains where alleles were sampled with equal probability from one of the six parents, and costs were sampled from a discrete uniform distribution, ranging from $20 to $200 in steps of $10. Strain picking, phenotype simulation, and power assessment was performed as above.

#### GWAS Phenotype Creation

We created sets of 100 simulated phenotypes each for our simulation studies. For each simulated phenotype, we drew a SNP at random that matched a required Minor Allele Frequency (MAF) criterion (5-10%, 10-30% or 40-50%). We then selected a β value for this SNP based on a desired χ^2^ threshold (10, 50 or 100) and added noise based on a multivariate normal (MVN) derived from two variance terms, the first the proportion of variance attributable to genetic effects 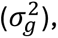 set at 40% consistent with our past work^56^ and the second the proportion attributable to all other sources of variation 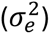 as follows:

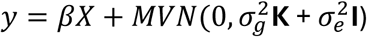

where **K** is the kinship matrix of the strains of the HMDP calculated using the kinship function of the myFastLMM R package^57^.

#### GWAS Approach

Simulated phenotype-genotype pairs were analyzed using the linear mixed model approach fastLMM^57^. Power was calculated based on the number of associations that met a genome-wide significance threshold of 4.2E-6 as previously determined in the HMDP^17,44,46,58^. At a χ^2^ of 100, there was 100% power in the full panel(Figure S1A). In the χ^2^ of 50 set, power ranged from 82-93% depending on the Minor Allele Frequency (MAF)(Figure S1A). Power in the χ^2^ of 10 set of simulated phenotypes was very low (<10%) in the whole panel and was not used going forward.

### Strain Subset Selection

We selected strain subsets for our simulation study using the following algorithms.

#### Greedy approaches

##### Random

Strains were selected randomly until the budget was exhausted.

##### MinCost

Strains were ranked from cheapest to most expensive, breaking ties randomly, then selected in that order until the budget was exhausted.

##### MaxMinNoCost

Strains were selected according to a greedy algorithm that chooses the most diverged strain at every step, regardless of cost. This is represented by the following pseudocode, where *n* is the total number of strains, *x* is a binary vector with *x_i_* representing whether strain *i* is selected, c*_i_* is the cost of strain *i*, *B* is the total budget, and *d_i,j_* is the genetic distance between strains *i* and j:

1. cheapest = which(c == min(c))
2. x[sample(cheapest, 1)] = 1 *# random initialization*
3. while (B > 0):

a. candidates = which(x == 0 & c <= B)
b. selected = which(x == 1)
c. for (a in candidates):

i. mindist_a = min(d_{a,s}) for s in selected)
d. new = which(mindist_a == max(mindist_a))[1]
e. x[new] = 1
f. B = B – c[new]

##### MaxMinMyopic

This method uses the same algorithm as MaxMinNoCost, except that the minimum distances are scaled by cost, meaning that step 3d would read:

- new = which(mindist_a/c_a == max(mindist_a/c_a))[1]

##### MaxSumNoCost

This method uses a similar approach to MaxMinNoCost, but steps 3c-d are different:

1. cheapest = which(c == min(c))
2. x[sample(cheapest, 1)] = 1
3. while (B > 0):

a. candidates = which(x == 0 & c <= B)
b. selected = which(x == 1)
c. for (a in candidates):

i. sumdist_a = sum(d_x, y for x, y in union(a, selected))
d. new = which(sumdist_a == max(sumdist_a))[1]
e. x[new] = 1
f. B = B – c[new]

##### MaxSumMyopic

As MaxSumNoCost, but as before, the sum of distances is scaled by cost in step 3d:

- new = which(sumdist_a/c_a == max(sumdist_a/c_a))[1]

##### MaxMAFNoCost

This method uses a similar framework as the algorithms above, where we choose the next strain such that it minimizes the following equation:

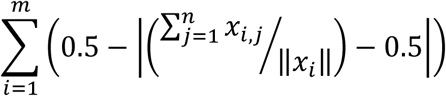

Here, x is a m x n matrix in which there are m SNPs; the first (n-1) columns are the currently selected strains and the nth column is the ‘applicant’ strain. This algorithm acts to select strains such that they push the minor allele frequency (MAF) of the selected strains as close to 50% as possible, thereby attaining ‘maximum MAF’. In pseudocode:

1. cheapest = which(c == min(c))
2. x[sample(cheapest, 1)] = 1
3. while (B > 0):

a. candidates = which(x == 0 & c < B)
b. selected = which(x == 1)
c. for (a in candidates):

i. sumMAFs_a = sum(0.5 – abs(rowMeans(snp_matrix[, union(selected, a)]) – 0.5))
d. new = which(sumMAFs_a == max(sum_MAFs_a))[1]
e. x[new] = 1
f. B = B – c[new]

##### MaxMAFMyopic

As MaxMafNoCost, but as before, sumMAFs_a is replaced by sumMAFs_a / c_a in step 3d:

- new = which(mindist_a/c_a == max(mindist_a/c_a))[1]

#### Other heuristic approaches

##### p-MedianNoCost

Because greedy search does not work well for the p-Median problem, we instead used the fast Partitioning Around (k-)Medoids method^55^ implemented in the pam function in the “cluster” package in R^59^. The PAM algorithm for p-Median problems involves making iterative improvements to the set of selected medoids, but unlike a pure greedy search, it allows medoids to be replaced. To handle the cost constraint, we ran PAM with k increasing from 2 to n-1, where n is the total number of strains. We kept solutions that were under budget, and of these kept the result that selected the most strains.

#### Optimization based approaches

##### p-MedianOptim

We formulated this problem to be similar to minimum-ADCL^33^ and k-medoids, with an additional constraint on total cost and with no predetermined number k of representatives. Thus, we were interested in picking k medoids or representative strains with minimum distance to the closest non-chosen strains for each representative, with k allowed to vary between one and the number of strains.

The p-MedianOptim method then minimizes the following sum:

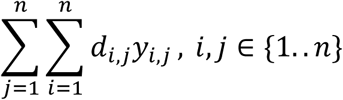

subject to the following constraints:

1. 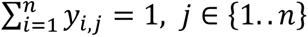
2. 0 ≤ *x*_*i*_ − *y*_*i*,*j*_ ≤ 1, *i*, *j* ∈ {1.. *n*}
3. *x*_*i*_, *y*_*i*,*j*_ ∈ {0, 1}, *i*, *j* ∈ {1.. *n*}
4. 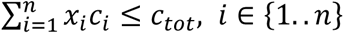
5. 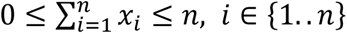

The first constraint guarantees that exactly one “nearest” representative is assigned to each of full set of strains. The second guarantees that *y*_*i*,*j*_is never 1 when *x*_*i*_ is not also 1: this in turn means that only distances from a representative to its assigned strains will be summed in the objective function. The third guarantees that *x* and *y* are binary, the fourth gives the cost constraint, and the fifth allows for any number of strains *k* (*k* = ∑*x*_*i*_) to be selected. Strains that must be selected (for example, strains that are already part of a researcher’s collection) can also be included as additional constraints on the value of a particular *x*_*i*_.

The method was implemented in R, using the “highs” package^60^ which provides bindings to the open-source HiGHS solver^61^.

We did not evaluate an ILP version of the MaxSum (or p-dispersion-sum) problem because even in tests on very small datasets, its performance was extremely slow. We also did not attempt to frame MaxMAF as an ILP problem because it uses the entire allele frequency matrix instead of a distance matrix, and would therefore require orders of magnitude more variables than the other ILP methods tested here.

### Fitting and Significance Testing

We fit generalized additive models (GAMs) to the power estimates, assuming the data were beta-distributed, using the restricted maximum-likelihood estimator the R package “mgcv”^62^. Because all methods eventually converge given a high enough budget, for the purpose of the model fit, we kept only budget values where the average power across every method was less than 90% of the maximum power observed. To reduce the impact of exact zeros at the low end, which distort the fit, we also slightly moderated power values by adding 0.005 and divided by 1.005. Power was then modeled as a per-method smooth function of the budget, using k=5 knots. After fitting GAMs, the estimated marginal means over these budget ranges were calculated using the “emmeans” R package. Using the “multcomp”^63^ package, significant pairwise differences at p=0.05 were then assessed using the Tukey adjustment, from which compact letter displays were finally generated.

### Validation Approaches

#### HMDP

The Right Ventricular weight phenotype after isoproterenol stimulation was selected as it had the largest number of genome-wide significant loci from its manuscript (Figure 3A)^18^. The mice used in the selected study did not comprise the entire panel, but instead consisted of 93 strains, which we calculated to be a total cost of $119,241. Costs were set at $20,000 intervals from $0 to $100,000 and all selection algorithms were run ten times per approach. Included SNPs were recalculated for each sample due to changes in MAF across the population due to removal of strains. Phenotypes were then subset and GWAS run through fastLMM^57^ followed by analysis of how many significant SNPs were maintained after subsetting as well as how many SNPs which had been present in the original panel were lost due to shifts in MAF.

## RESULTS

### Bacterial Phylogenetic Regression Studies

We considered two scenarios for our test of strain selection methods on a bacterial dataset. In the first of these, we used the *Lachnospiraceae* phylogenetic tree and genotypes that we inferred from the genomes, with genotype represented as presence or absence of “phylogroups” (sets of orthologs that had identical phylogenetic profiles). We then simulated phenotypes correlated with a randomly selected phylogroup, such that a phylogenetic regression on the full panel would have approximately 80% power to detect an effect at a nominal p-value ≤ 0.05 (Figure 2).

**Figure 2.**
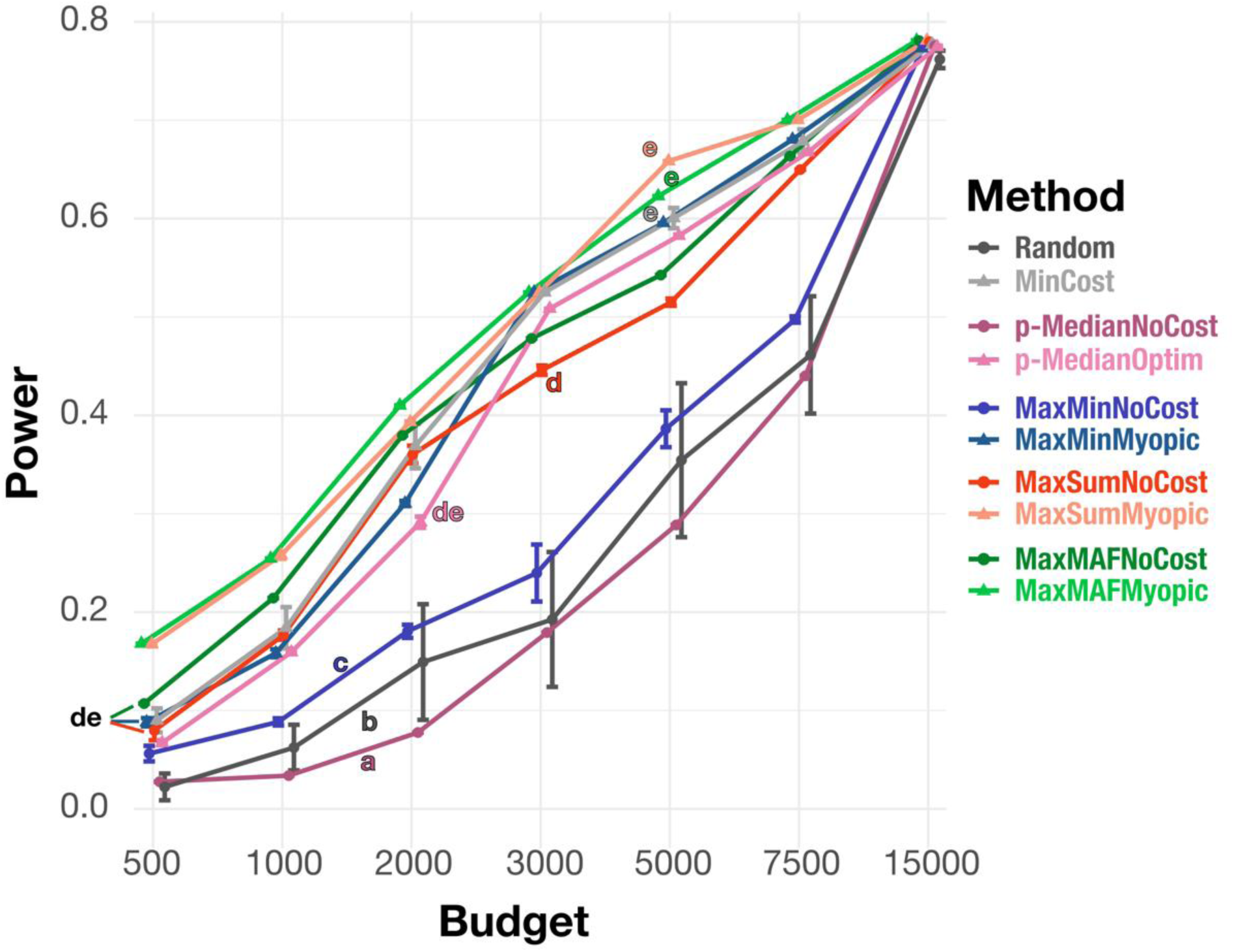
Power of phylogenetic regressions on traits simulated across 165 commercially available *Lachnospiraceae* isolate genomes. Methods are colored by how they consider genetic diversity (none: gray; p-Median: purple; MaxMin: blue; MaxSum: red; MaxMAF: green), with lighter colors corresponding to methods that consider cost (MinCost and the Myopic and Optim methods). Error bars are ±1 SD of the mean of five rounds of picking genomes. Lowercase letters (a-d) indicate significantly different groups; methods are significantly different at p=0.05 if they do not share a letter (GAM beta regression, Tukey-adjusted post-hoc pairwise tests; see Methods).

Selecting strains randomly, as expected (Random), fared poorly in terms of power. More interestingly, though, despite the amount of prior literature focusing on these methods, we found that the greedy MaxMin method that did not consider costs was barely better than random guessing, and the heuristic for the p-Median problem without costs was actually even worse. We believe this poor performance can be explained by the fact that costs in our panel varied over more than an order of magnitude (from $37.33 to $474.00), and so methods that did not consider costs explicitly tended to select far fewer strains than their cost-aware counterparts (Table 1).

**Table 1:** Median number of *Lachnospiraceae* strains selected and range (in brackets), by method and budget (“Real” simulation).

| Budget | 500 | 1000 | 2000 | 3000 | 5000 | 7500 |
| --- | --- | --- | --- | --- | --- | --- |
| Random | 4.6 [1-7] | 10.4 [6-14] | 22 [15-28] | 29.2 [23-36] | 54.8 [40-62] | 74.8 [64-86] |
| p-MedianNoCost | 5 [5-5] | 6 [6-6] | 20 [20-20] | 34 [34-34] | 54 [54-54] | 84 [84-84] |
| MaxMinNoCost | 8.2 [7-9] | 14 [14-14] | 27.6 [26-28] | 36.2 [33-40] | 64.8 [60-68] | 90.2 [89-92] |
| MaxSumNoCost | 9 [9-9] | 14.4 [14-16] | 26.4 [26-28] | 39.2 [38-40] | 53.8 [53-55] | 94 [94-94] |
| MaxMAFNoCost | 9 [9-9] | 14 [14-14] | 26 [26-26] | 40 [40-40] | 61 [61-61] | 93 [93-93] |
| p-MedianOptim | 13 [13-13] | 26 [26-26] | 51 [51-51] | 78 [78-78] | 97 [97-97] | 119 [119-119] |
| MaxMinMyopic | 13 [13-13] | 26 [26-26] | 53 [53-53] | 79 [79-79] | 97 [97-97] | 119 [119-119] |
| MaxSumMyopic | 13 [13-13] | 26 [26-26] | 53 [53-53] | 79 [79-79] | 97 [97-97] | 119 [119-119] |
| MaxMAFMyopic | 13 [13-13] | 26 [26-26] | 53 [53-53] | 79 [79-79] | 98 [98-98] | 119 [119-119] |
| MinCost | 13 [13-13] | 26 [26-26] | 53 [53-53] | 79 [79-79] | 98 [98-98] | 119 [119-119] |

Even the cost-aware versions of MaxMin or p-Median, however, were not the strongest performers overall. In fact, even the non-cost-aware MaxSum method, which maximizes the total sum of pairwise distances, and the MaxMAF method, which maximizes the total sum of minor allele frequencies, had performance that approached the cost-weighted version of MaxMin and p-Median. The cost-aware versions of MaxMAF and MaxSum preserved even more power, and indeed, were essentially tied for the highest performance overall. This difference was especially pronounced at lower budgets (e.g., $500-$1,000).

We were surprised to see that one of the simplest methods — picking strains in order of cost, without considering diversity at all (MinCost) — was the third-ranked, behind MaxMAFMyopic and MaxSumMyopic, and was not significantly worse than these across the range of budgets. MinCost even beat the globally-optimized p-MedianOptim method in these simulations (though not significantly). As above, a possible explanation is that MinCost always maximizes the number of strains selected (Table 1), which, all else being equal, should always increase power.

We reasoned that we might expect to see MinCost perform worse in settings where the panel contained clusters of inexpensive but closely related strains, as these would make only marginal contributions to the diversity metrics we consider. In some collections, for example, certain individual species with high value in the wet-lab or industry, such as *E. coli* or *B. subtilis*, have been “oversampled” relative to more distant phylogenetic neighbors. We would not have expected to observe this phenomenon in the *Lachnospiraceae* data set we considered above because this set does not contain common model or industrial microbes (at least, not yet). To test this hypothesis, we therefore simulated the addition of another collection that had a large number of inexpensive, closely-related strains, which we refer to as “oversampled” strains below.

In this “oversampled” scenario, we indeed saw a much larger advantage for the methods that consider both cost and diversity, and also saw more differentiation among these methods. Strikingly, MaxMAFMyopic still performed the best overall, significantly outperforming MinCost across all budgets below the very highest (where all methods performed well). In this setting, the p-MedianOptim method was second-best overall. Because p-MedianOptim minimizes the distance of non-selected to selected strains, it should pick strains that are evenly distributed in terms of their genetic relatedness (see Figure 1), which may explain why it is the most robust to oversampling a particular taxon. The next best methods were, in order, MaxSumMyopic and MaxMinMyopic. All four of these methods significantly outperformed MinCost across the range of budgets, again matching the intuition that prioritizing diversity becomes more important when a low-diversity clade is oversampled.

Unexpectedly, MaxSumNoCost and MaxMinNoCost were significantly worse than random guessing. Because methods that considered diversity in this experiment tended to perform so well, this was initially surprising. However, in this experiment, the highly-similar strains we simulated were also very low cost ($20.00). This gives the Random approach an advantage, as it would tend to run out of its budget slower than the other NoCost methods, which are required to maximize diversity. Indeed, MaxMinNoCost and MaxSumNoCost picked the fewest total genomes out of any method (Table 2). We found there was no consistent relationship between performance and the number of synthetic “oversampled” strains that were selected (Table 3). MaxMinNoCost and MaxMAFNoCost picked the least but performed worst, while at low budgets (≤$5000), p-MedianOptim selected more than Random and MaxMinMyopic.

**Table 2:** Median number of total strains selected and range (in brackets), by method and budget (“oversampled” simulation).

| Budget | 500 | 1000 | 2000 | 3000 | 5000 | 7500 |
| --- | --- | --- | --- | --- | --- | --- |
| Random | 8.8 [5-11] | 12.4 [10-18] | 27.2 [24-36] | 38.4 [28-47] | 74 [67-82] | 104.8 [85-121] |
| p-MedianNoCost | 8 [8-8] | 15 [15-15] | 31 [31-31] | 41 [41-41] | 59 [59-59] | 74 [74-74] |
| MaxMinNoCost | 8.6 [7-9] | 14 [14-14] | 22.8 [21-24] | 28 [28-28] | 48.4 [47-52] | 65.2 [64-66] |
| MaxSumNoCost | 7 [7-7] | 12.4 [12-14] | 19 [19-19] | 30 [28-32] | 44.6 [43-45] | 74 [74-74] |
| MaxMAFNoCost | 7 [7-7] | 12 [12-12] | 26 [26-26] | 38 [38-38] | 63 [63-63] | 93 [93-93] |
| p-MedianOptim | 16 [16-16] | 30.4 [30-31] | 58 [58-58] | 85 [85-85] | 117 [117-117] | 141 [141-141] |
| MaxMinOptim | 14 [14-14] | 27 [27-27] | 54 [54-54] | 78 [78-78] | 97 [97-97] | 119 [119-119] |
| MaxSumMyopic | 17 [17-17] | 34 [34-34] | 68 [68-68] | 100 [100-100] | 165 [165-165] | 198 [198-198] |
| MaxMAFMyopic | 14 [14-14] | 27 [27-27] | 54 [54-54] | 81 [81-81] | 102 [102-102] | 121 [121-121] |
| MinCost | 20 [20-20] | 40 [40-40] | 80 [80-80] | 113 [113-113] | 166 [166-166] | 198 [198-198] |

**Table 3:** Median number of the additional oversampled strains that were selected and range (in brackets), by method and budget (“oversampled” simulation).

| **Budget** | **500** | **1000** | **2000** | **3000** | **5000** | **7500** |
| --- | --- | --- | --- | --- | --- | --- |
| **Random** | 4.6 [2-7] | 5.4 [3-9] | 11.4 [8-13] | 13.2 [8-17] | 27.2 [23-31] | 39.2 [28-49] |
| **p-MedianNoCost** | 5 [5-5] | 5 [5-5] | 7 [7-7] | 8 [8-8] | 10 [10-10] | 10 [10-10] |
| **MaxMinNoCost** | 1.2 [1-2] | 1.6 [1-2] | 1.4 [1-2] | 2 [2-2] | 1.8 [1-2] | 2 [2-2] |
| **MaxSumNoCost** | 1.2 [1-2] | 1.4 [1-2] | 2 [2-2] | 2.2 [2-3] | 1.8 [1-2] | 2 [2-2] |
| **MaxMAFNoCost** | 2 [2-2] | 2 [2-2] | 4 [4-4] | 5 [5-5] | 10 [10-10] | 16 [16-16] |
| **p-MedianOptim** | 8 [8-8] | 11.2 [10-13] | 14 [14-14] | 15 [15-15] | 27 [27-27] | 29 [29-29] |
| **MaxMinMyopic** | 2 [2-2] | 2 [2-2] | 2 [2-2] | 8 [8-8] | 21 [21-21] | 20 [20-20] |
| **MaxSumMyopic** | 12 [12-12] | 23 [23-23] | 44 [44-44] | 61 [61-61] | 95 [95-95] | 100 [100-100] |
| **MaxMAFMyopic** | 2 [2-2] | 1 [1-1] | 2 [2-2] | 2 [2-2] | 5 [5-5] | 2 [2-2] |
| **MinCost** | 20 [20-20] | 40 [40-40] | 80 [80-80] | 100 [100-100] | 100 [100-100] | 100 [100-100] |

Overall, failing to consider costs appears to harm performance more than failing to consider diversity. In the presence of strongly unequal costs, maximizing diversity alone can actually be worse than random guessing. However, the best-performing methods consider both. The cost-aware MaxSum and MaxMAF methods were the most powerful overall, and were joined by the cost-aware p-Median method when the panel was expanded to include many closely-related strains.

### Mouse Genome Wide Association Studies

We next evaluated these methods using simulations conducted in a substantially different context: genome-wide association studies (GWAS) on subsets of the Hybrid Mouse Diversity Panel (HMDP), which consists of approximately 175 strains of mice. At our selected effect size (χ^2^) and minor allele frequency (MAF) thresholds, we observe approximately 50% power to detect a true positive result at ∼60 strains at a χ^2^ of 100 and a MAF of 10-30%, or 100-110 strains at a χ^2^ of 50 at various MAF thresholds (Fig S1A). At χ^2^=100, there is 100% power to detect phenotypes in the whole panel, while at χ^2^=50, there is 82-93% power to detect a phenotype.

With this understanding of how random strain selection is likely to affect GWAS results in our panel, we set out to examine the behavior of each of selection approaches at price limits ranging from $20k to $240k (approximately the cost of the entire panel). We observe some patterns which are consistent across all χ^2^ and MAF parameters (Fig. 4). The first is a very rapid jump in power in some models at low cost thresholds (see, for example, the jump from 10 to 50% power in Figure 4D from $20k to $40k). This reflects the specific cost structure of these strains: there are a large number of relatively inexpensive mice that can all be obtained at $40k total budget, while many of strains found in the Recombinant Inbred panels are significantly more expensive.

Several results line up with our observations on the bacterial lines. The Random, MaxSumNoCost, and MaxMinNoCost algorithms all performed significantly worse (Tukey-adjusted p<0.05) than their cost-aware Myopic counterparts. Typically, MinCost also remained in a statistical dead heat for the top rank.

However, some methods showed reversals from our bacterial simulations. We observed that MaxMinNoCost, instead of p-MedianNoCost, was either the worst or second-worst performer at all cost/χ^2^/MAF combinations, and in fact, p-MedianNoCost was actually tied for the top place. Most surprisingly, MaxMAFMyopic, previously in the top two methods, performed worse than average. In fact, MaxMAFMyopic was often beaten significantly not only by MaxSumNoCost, but by its own cost-unaware counterpart, MaxMAFNoCost (Fig. 4).

We sought to understand these differences in performance by first testing the impact of two parameters that might differ across settings: first, effect size (χ^2^), and second, the minor allele frequencies (MAFs) of the causal SNPs (Fig. 4A-C). We found that effect size appeared to mainly determine how much power we retain overall by subsampling, without large method-specific differences (compare Fig. 4A and 4C). As expected, for very strong signals (χ^2^ =100; Fig. 4A), power rose faster with the available budget, with the best methods reaching 90% power at a $100K budget, compared to less than 50% for weaker signals (χ^2^ =50; Fig. 4C). However, in both cases, methods clustered into approximately the same three groups: the best were MaxSumMyopic, p-MedianOptim, p-MedianNoCost, and MinCost; a second tranche included both versions of MaxMAF, MaxMinMyopic and MaxSumMyopic; and the worst were Random and MaxMinNoCost.

In contrast, certain methods did show MAF-dependent behavior. We assessed this by sampling three different MAF cutoffs at an effect size of χ^2^ = 50: 5-10%, 10-30%, and 40-50% (Fig. 4B-D). We hypothesized that for low MAFs, methods that prioritize rare variants should maximize power. This is because low MAFs mean that the causal SNPs are rare, and because χ^2^ was held constant, it then follows that individual rare SNPs must have larger effects on the phenotype.

Indeed, we observed that MaxMAFNoCost, which would favor adding strains with rare variants, was the method affected most dramatically by allele frequency. When causal SNPs already have high MAFs (40-50%), prioritizing minor allele recovery should be unnecessary, as many strains contain the relevant mutant allele; and, because it does not consider costs, MaxMAFNoCost will lose power because it also selects fewer strains than the cost aware models (Fig. S1B). Sure enough, at high MAFs, MaxMAFNoCost performs poorly, at times worse than random selection (Fig. 4D). At moderate MAF (10-30%), MaxMAFNoCost and MaxMAFMyopic perform similarly and in the middle of the set of models in terms of power recovery, comfortably above Random, but significantly below MinCost (Fig. 4C). At low MAFs (5-10%), however, MaxMAFNoCost actually becomes the best model for budgets between $100k and $180k, at times recovering 10% more power than the second best model (Fig. 4B). This is likely because rare SNPs carry most of the signal in the data, and so while MaxMAFNoCost picks fewer strains (a bigger problem with very low budgets), it also tends to pick the ‘right’ strains for recovering the signal.

While we have previously only considered the cost of acquiring the strains, in most real-world scenarios, downstream phenotyping experiments require additional costs that scale per animal or strain, placing further economic constraints on how many strains it is feasible to purchase. To simulate these additional costs (which could include RNAseq, histology, experimental manipulation, etc.), we added $2,500 to the cost of each mouse strain. This also smooths out the very large cost differences seen in the mouse data, such that now the most expensive strain would only be 2x that of the least expensive strain, rather than >40x. Indeed, we see smaller differences in the number of mouse strains chosen between the models (Fig S1C).

With additional costs, MaxMAFNoCost’s advantage on low MAF SNPs widened even more across the entire budget range (Fig. 5A), at times beating the next best model by 25%. Furthermore, it became the top-ranked method at moderate MAFs as well (Fig. 5B). As we discussed above, MaxMAFNoCost is more likely to pick more ‘useful’ strains, but also fewer, leading to a power tradeoff. When strain costs become more similar, this second factor should become less important, which explains the improvement in power across MAF thresholds. Crucially, however, MaxMAFNoCost still performed poorly when MAFs were between 40-50% (Fig. 5C). Thus, maximizing MAF without regard to cost is a viable strategy when causal alleles have low frequency, and especially as cost differentials decrease, but not when MAFs are already close to parity.

With less differentiation by cost, the other competing methods became more similar to one another (Fig. 5A-B), except for MaxMinNoCost, which remained weaker than random guessing. We also did not observe the same early jump in power we saw previously, which was expected as there was less of a cost differential between the cheapest and most expensive strains. Interestingly, MaxMAFMyopic’s performance became more variable, especially at low MAF, but still lagged behind the others. MinCost, in contrast, remained statistically tied for the best or second-best method across these simulations.

We wished to further explore the counterintuitive result that while optimizing for either cost (MinCost) or total MAF (MaxMAFNoCost) separately was effective for picking mouse strains, optimizing an objective that included both factors was not, even though this was the best approach for bacteria. We entertained two explanations. First, there could be a bias where more expensive mouse strains contain rarer alleles, meaning that the strains that maximize MAF were intrinsically high-cost. Second, it could be that the greedy MaxMAFMyopic algorithm simply did not get close enough to a global optimum on the mouse data. This failure could be because the genetic architecture of the two groups differs: the mouse genomes include recombinant inbred lines, while the bacteria are natural isolates from many different species.

We tested the first of these hypotheses by randomly distributing costs (pulling from a uniform distribution from $60 to $1000 per strain) and re-running the simulations. When we did so, we still observed that MaxMAFNoCost preserves more power than MaxMAFMyopic (Tukey-corrected p<0.05, Figure S4), indicating that a correlation between rare alleles and expensive strains does not drive this pattern. To test the second explanation, we took a set of bacterial genomes and generated simulated “recombinant” strains, whose genotype at each locus was randomly sampled from the “parental” strains, and then repeated the analysis. We found that in this experiment, all of the cost-aware approaches performed similarly, outperforming the non-cost-aware methods, with the sole exception of MaxMAFMyopic (Figure S5). Together, these two results favor the explanation that a simple myopic method may not effectively co-optimize MAF and cost when the full panel contains a large number of recombinant lines, which largely contain different permutations of the same variants, as opposed to isolates, which have more unique variants per strain.

### GWAS Validation in the HMDP

Finally, we applied our selection criteria to real data pulled from a prior HMDP study on cardiac dysfunction^18^. In particular, we examined the effects of strain selection on GWAS loci for right ventricle weight after catecholamine challenge, which had the largest number of genome-wide significant SNPs in the original study (Fig 3A).

**Figure 3:**
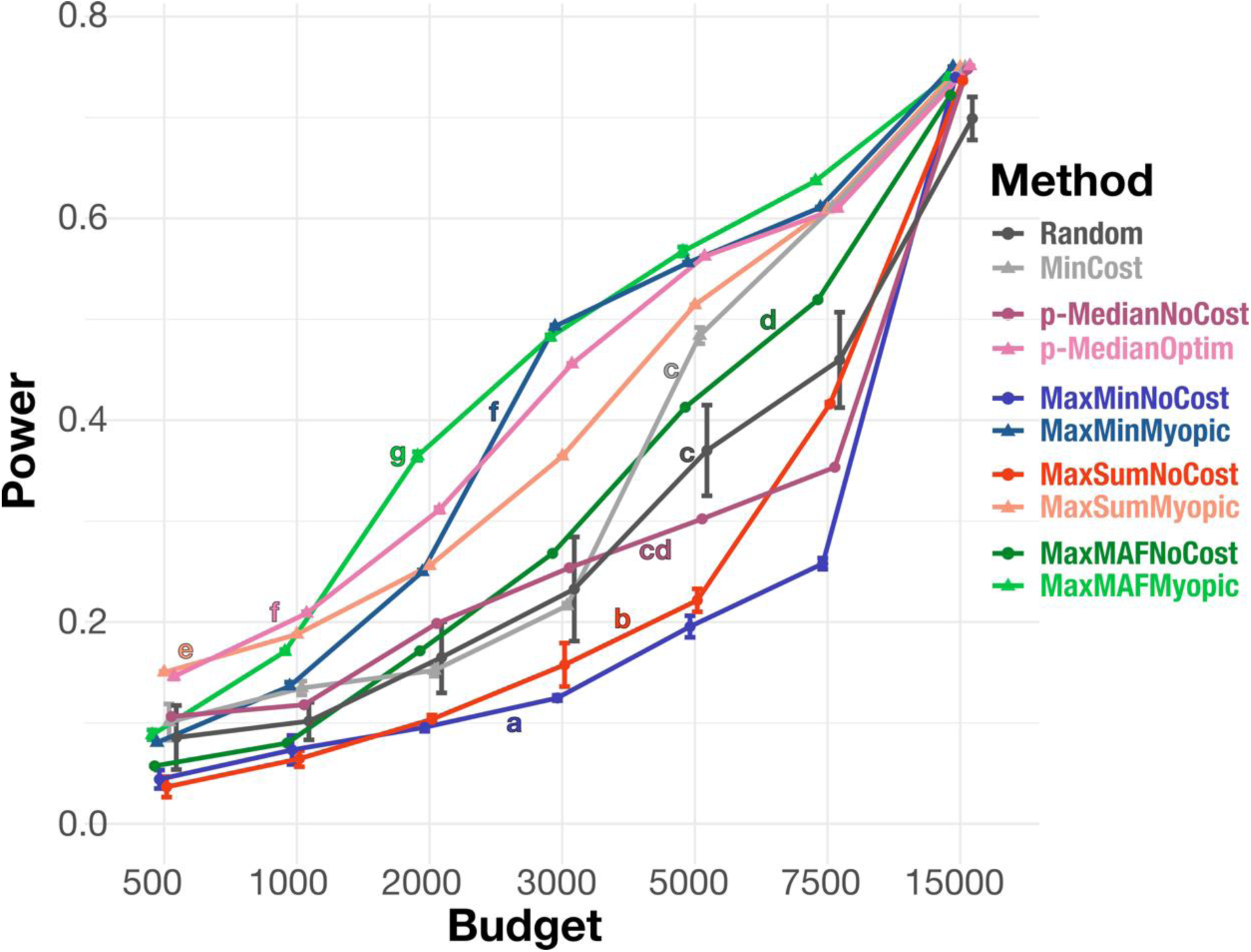
Power of phylogenetic regressions on traits simulated across 165 commercially available *Lachnospiraceae* isolate genomes, augmented by 100 highly similar synthetic “oversampled” strains. Methods are colored as in Figure 2. Error bars are ±1 SD of the mean of five rounds of picking genomes. Lowercase letters (a-d) indicate significantly different groups; methods are significantly different at p=0.05 if they do not share a letter (GAM beta regression, Tukey-adjusted post-hoc pairwise tests; see Methods).

To be consistent with the original study, SNPs with a MAF of less than 5% were excluded; because strain selection affects this calculation, we first examined the effects of removing strains on the number of SNPs tested. As expected from our simulations, MaxMinNoCost and Random resulted in the largest number of SNPs being lost, over 10% at $20K and 5% at $60K. MaxMAFNoCost predictably lost the fewest number of SNPs, <3% at $20K and only 0.3% at 60K (Fig 6B). MinCost, despite not considering genetic diversity at all, did reasonably well, with 4.3% SNP loss at 20K and 1.8% loss at 60K.

We next queried how many genome-wide significant SNPs were identified for each method at each cutoff when compared to the full panel (Fig 6C). Strikingly, at $80K and $100K cutoffs, MaxSumMyopic actually identified more genome-wide significant loci than the full panel, joined by MaxMinMyopic at the highest threshold. Random selection resulted in very high variance, with some permutations returning as many as 433 significant SNPs (compared to 157 found in the real data), while others at the same $60K threshold returned as few as 27. Consistent with our simulations, MaxMinNoCost recovered very few significant SNPs, returning zero significant hits at every budget other than the highest. We were surprised that the two MaxMAF algorithms also performed poorly; however, when we looked at the significant SNPs from the real panel, we found that many of the significant SNPs had MAFs >20%, a setting where MaxMAF performed less well in simulations (Fig. 4C-D).

**Figure 4:**
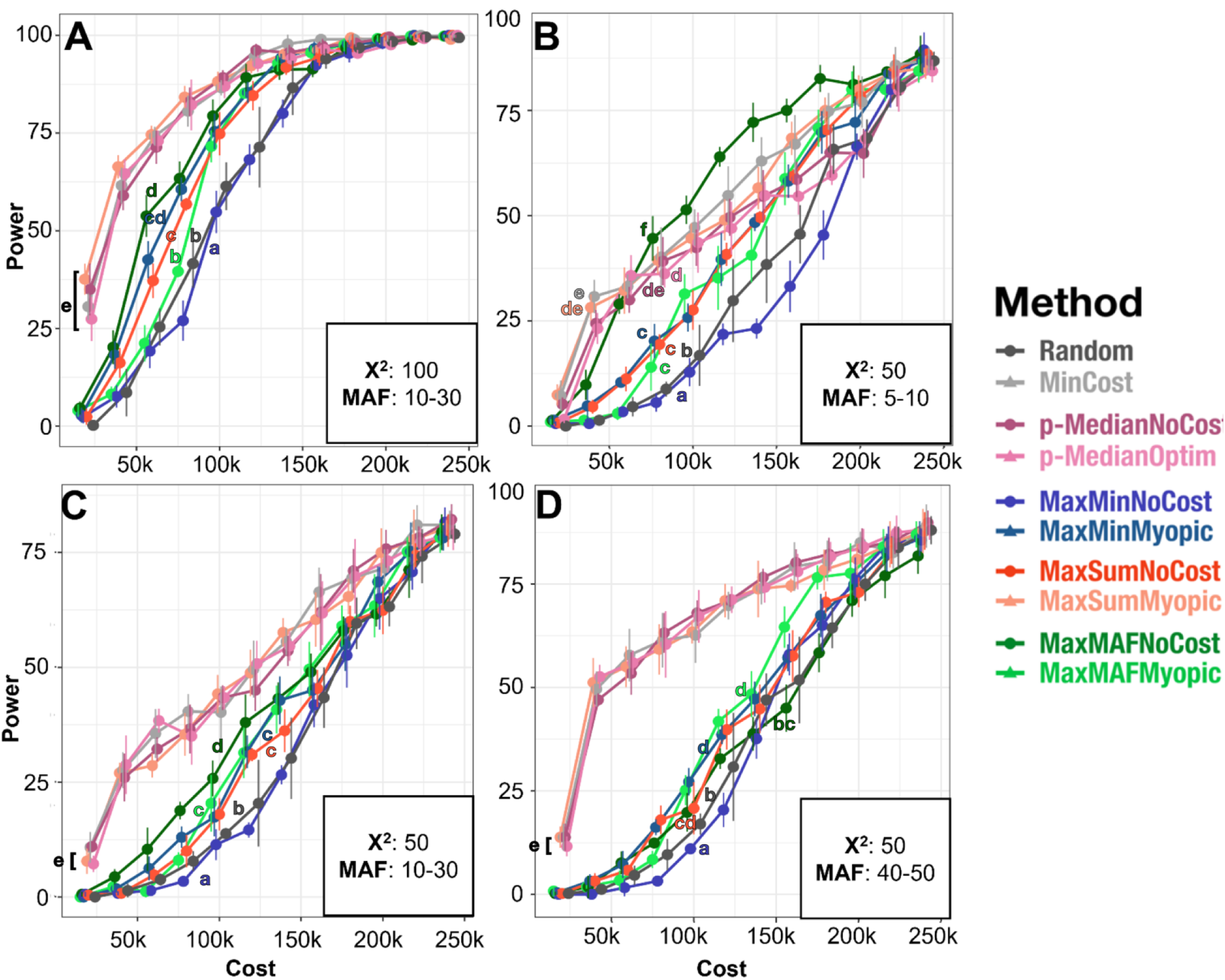
Mouse GWAS Power Simulations Without Additional Costs Per Strain. Results of simulation studies at $20k intervals from 20-240k. **A)** χ^2^ of 100 and MAF of 10-30%, **B)** χ^2^ of 50 and MAF of 5-10%, **C)** χ^2^ of 50 and MAF of 10-30%, **D)** χ^2^ of 50 and MAF of 40-50%. N=100 simulated phenotypes for each of 5 replications. Values are slightly jittered on the x axis to allow for observation of each method’s error. Error bars are ±1 SD. Lowercase letters (a-f) indicate significantly different groups; methods are significantly different within a panel at p=0.05 if they do not share a letter (GAM beta regression, Tukey-adjusted post-hoc pairwise tests; see Methods).

Finally, we tested agreement with the results from the real panel. We first asked what percentage of the significant SNPs from the real panel were recovered in our sub-panels (Fig 6D); assuming that the SNPs from the real panel are valid, this is a measure of recall. Here, p-MedianOptim and MinCost performed the best, recovering about 37% of all SNPs at $60K and, in the case of p-MedianOptim, 75% at 80K. Notably, despite returning the most overall SNPs, with comparable numbers to the full panel starting at $40K and exceeding it at $80K, MaxSumMyopic did not recover a majority of the SNPs from the real panel until reaching $100K, suggesting an elevated false positive rate.

We next calculated the percentage of SNPs recovered by the sub-panels that were also found in the real panel (Figure 6E), a measure of precision. Indeed, MaxSumMyopic had fairly low precision compared to other methods. Conversely, MaxMAFNoCost, despite returning very few significant SNPs, was very faithful to the full panel, with 100% of its recovered SNPs being reproduced (other than at $100k, where it drops to 98%). Again, the best methods were p-MedianOptim, followed by MinCost, with rates between 45-80% starting at $40K onward. Thus, p-MedianOptim and MinCost optimized both precision and recall the best, while the selections made by MaxMAFNoCost strongly favored precision.

## DISCUSSION

In this study, we set out to address a common problem faced by researchers: how best to select strains from an available panel in order to stay within a fixed budget while retaining the power needed to make reliable discoveries. Because such comparative approaches are used in many different fields of study within biology, we simulated experiments on phylogenetic regression in microbes and on genome wide association studies in mice, followed by subsampling of real mouse GWAS data as a validation.

We find that in both of these settings, and under most conditions we tested, explicitly accounting for cost mattered as much as accounting for diversity. In realistically-priced strain collections, methods that ignored costs selected fewer, if potentially more diverse, strains, driving down power. Conversely, simply maximizing sample size, by selecting strains from cheapest to most expensive until the budget is exhausted (MinCost), was surprisingly competitive compared to models that combined both cost and diversity considerations. Typically, the highest-ranked methods considered both cost and diversity, though which specific diversity-maximizing objective performed best depended on the setting, and in many cases, MinCost statistically tied these methods. Surprisingly, even though methods related to MaxMin have been explored much more in the genetics literature, we found that MaxMin performed poorly, and that the best diversity objectives were usually MaxSum or MaxMAF, and in some cases p-Median. We discuss our results in more detail below, along with a few notable exceptions to these trends.

### Why does MinCost perform better than expected?

We did not expect MinCost to outperform many methods that explicitly maximize diversity. However, while MinCost itself does not consider genetic distance, this does not mean diversity is irrelevant. Researchers who contribute to collections are likely motivated, at least in part, by considerations like expanding the diversity of cultured microbes to which we have access, or in the case of mice, breeding lines that aim to randomly shuffle genotypes through recombination. In the context of a panel that is already diverse, “buy as many as you can afford” becomes a viable strategy because it maximizes sample size, and therefore the chance to sample informative genotype/phenotype pairings. Indeed, when we changed this picture by introducing a large number of cheap, closely related bacterial strains, we saw that MinCost’s efficacy began to falter, and considering genetic diversity became increasingly important. While methods like p-MedianOptim, and, to a lesser extent, MaxMinMyopic were tied with or worse than MinCost on the original data, in an oversampled panel they outperformed MinCost at many budget thresholds.

### Why are MaxMin methods outperformed by alternatives?

We found that MaxSum and MaxMAF, and to a lesser extent p-Median, commonly outperformed the more commonly-studied MaxMin method. One reason for MaxSum’s superior performance may be that MaxMin penalizes potential additions for being closely-related to even one selected strain. In contrast, the MaxSum method maximizes the sum of all pairwise distances. This means that being closely-related to one strain can be balanced out by being very diverged from others in the set. Thus, MaxSum may be less likely to reject potentially informative strains that would be deprioritized by MaxMin.

Intriguingly, MaxMAF showed even stronger performance across simulations than MaxSum. We explain this by noting that the power of a regression analysis is typically higher when the independent variable has a larger sample variance, and binomial distributions have the highest variance at a mean of 0.5. Because MaxMAF pushes the average MAF towards 0.5, it is therefore selecting for larger sample variances, which MaxMin does not consider.

Allele frequency has, in some ways, a more direct connection to genome-wide association studies than measures of diversity based on genetic distances, as GWAS relies on the recovery of information from individual informative SNPs or CNVs. In contrast, for example, bioconservation studies often aim to maximize the total expected functional diversity in a set of taxa, which may be more closely related to the overall genetic distance between them. When causal allele frequencies are high, these two approaches may give similar answers, but at low frequencies, ensuring more rare alleles have sufficient variation may have a greater impact than maximizing genetic distances among selected taxa.

### How does genetic architecture affect strain selection?

The two settings we tested represent different types of genetic diversity: cross-species variation in bacterial gene family content, and within-species variation in mouse genetic polymorphisms. We expect a high degree of nested structure in the bacterial genotypes because many genes should be correlated to phylogeny and some may in fact be clade-specific. In contrast, the mouse data includes many recombinant inbred lines (RILs). Unlike the bacterial isolates, these were bred specifically with the aim of decorrelating genetic variants from one another and thus facilitating trait mapping. In our dataset, bacterial genes also tended to have much lower MAF on average than mouse SNPs. Comparing these two settings can therefore give us a window into how the genetic architecture of strains and/or traits can affect the performance of strain selection methods.

Certain trends were consistent: *most* methods that ignored prices (Random, MaxMinNoCost, MaxSumNoCost, p-MedianNoCost) underperformed their cost-aware counterparts in both bacteria and mice. MaxMAF was an exception, with the NoCost version typically performing better than Myopic; as noted above, however, the Myopic version also performed poorly when we simulate recombinant bacterial progeny from a small set of parental strains, suggesting that this reversal is actually a failure of optimization.

While changing the effect size did not change the relative rankings of most approaches, we did observe an effect of causal allele frequency, especially for MaxMAFNoCost and also for MaxMAFMyopic. MaxMAFNoCost’s single-minded focus on driving allele frequencies towards 50% leads to the lowest number of selected strains of any approach (Figure S1B), which can harm power when the causal allele is high MAF. However, as noted above, because MaxMAF retains more rare alleles, it becomes one of the best algorithms when traits are driven by rare SNPs, besting all others in the $160K-$240K budget window (Fig. 5A). This also partially explains why the MaxMAF methods performed so well in bacteria, where the distribution of MAFs is shifted lower (compare, e.g., Fig. S2A, B, and D).

**Figure 5:**
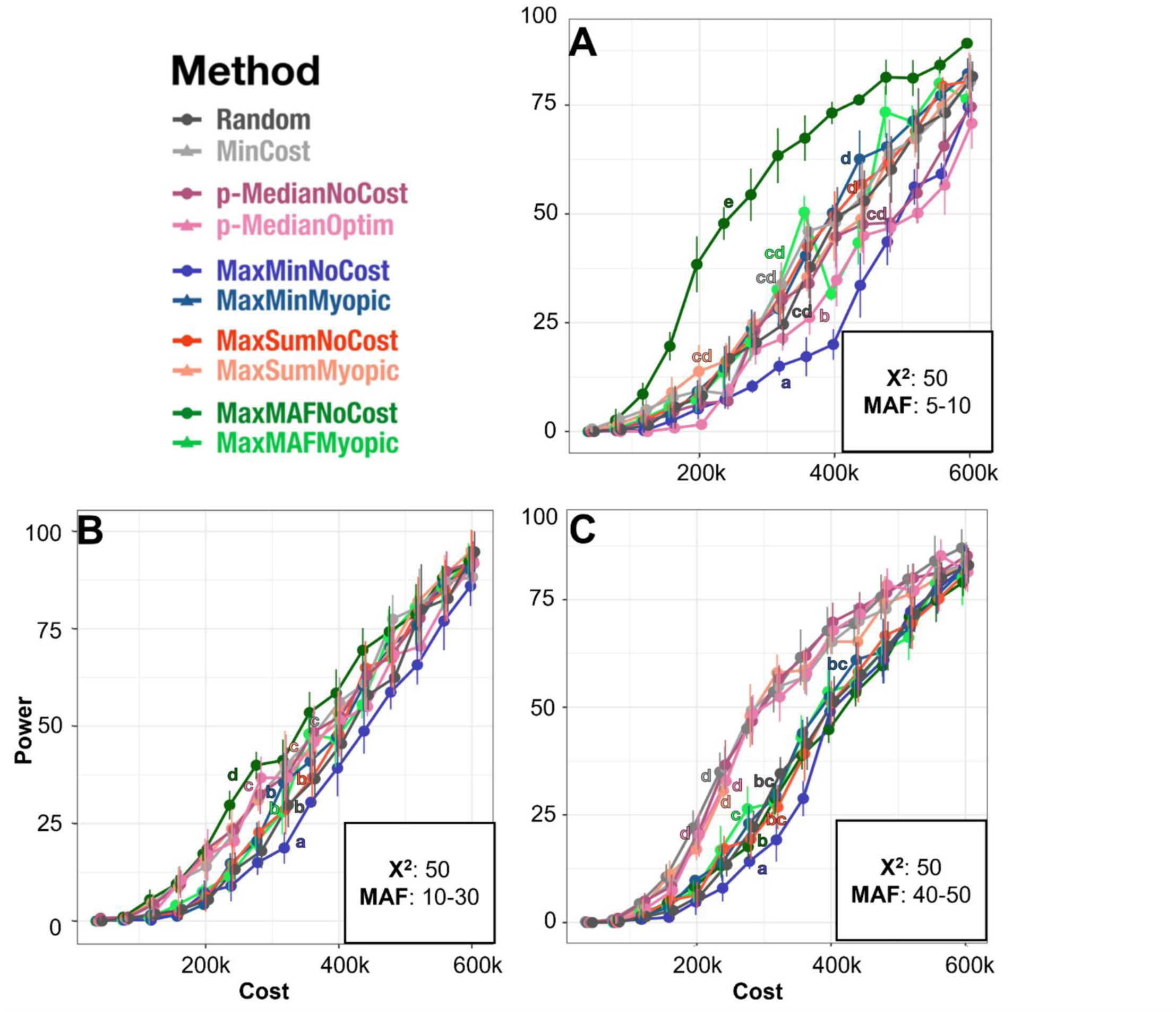
Mouse GWAS Power Simulations With Additional Cost Per Strain. Results of simulation studies after $2,500 was added to the cost of each strain at $40k intervals from 40-600k. **A)** χ^2^ of 50 and MAF of 5-10%, **B)** χ^2^ of 50 and MAF of 10-30%, **C)** χ^2^ of 50 and MAF of 40-50%. N=100 simulated phenotypes for each of 5 replications. Values are slightly jittered on the x axis to allow for observation of each method’s error. Lowercase letters (a-f) indicate significantly different groups; methods are significantly different within a panel at p=0.05 if they do not share a letter (GAM beta regression, Tukey-adjusted post-hoc pairwise tests; see Methods).

### How does cost structure affect selection?

Strain prices can vary over orders of magnitude for both bacteria and mice. For example, a breeding trio for the very popular C57BL/6J strain costs less than $66 at time of writing, versus up to $2,850 for a litter of one of many cryopreserved lines^48^, a 47-fold difference. However, the expenses associated with downstream phenotyping experiments are typically large and constant across strains, partially equalizing the total cost.

We therefore further investigated the impact of adding additional costs to the mouse strains. This compressed a 47-fold difference between the least- and most-expensive strains to just 2.1-fold. One impact was on the shape of the power curve: in the original simulations, we saw a pronounced jump in power for the cost-aware methods between $20K and $40K (Fig. 4A, 4C), as these methods could take advantage of a large number of low-cost mouse strains (Figure S1B). After adding additional costs, though, the cost-aware versions no longer showed this early jump (Figure S1C), and the power increase was instead diffused across $40K-$280K (Figure 5). The most striking impact, however, was again seen with MaxMAFNoCost, which widened its lead at low MAF and became the top-ranked method at medium MAF (10-30%). This is likely because one of the main disadvantages of MaxMAFNoCost is that it tends to select the fewest strains; with less differentiation in strain costs, this disadvantage is attenuated relative to other methods (Fig. S1B-C).

### Can these methods recapitulate the results of a real GWAS study?

Applying our selection algorithms to a real HMDP GWAS (for right ventricular weight after catecholamine challenge, Figure 6A) on 104 strains of the HMDP confirmed many of our simulation results. Most methods which ignored costs (Random, MaxMinNoCost) lost many SNPs from the analysis set due to shifting MAFs dropping below the minimum 5% threshold used in the original study (Figure 6B), especially at low budgets. Again, the exception was MaxMAFNoCost, which actually preserved the most SNPs, losing less than 1% from our second lowest budget threshold onwards (compared to over 10% for MaxMinNoCost at this threshold).

**Figure 6:**
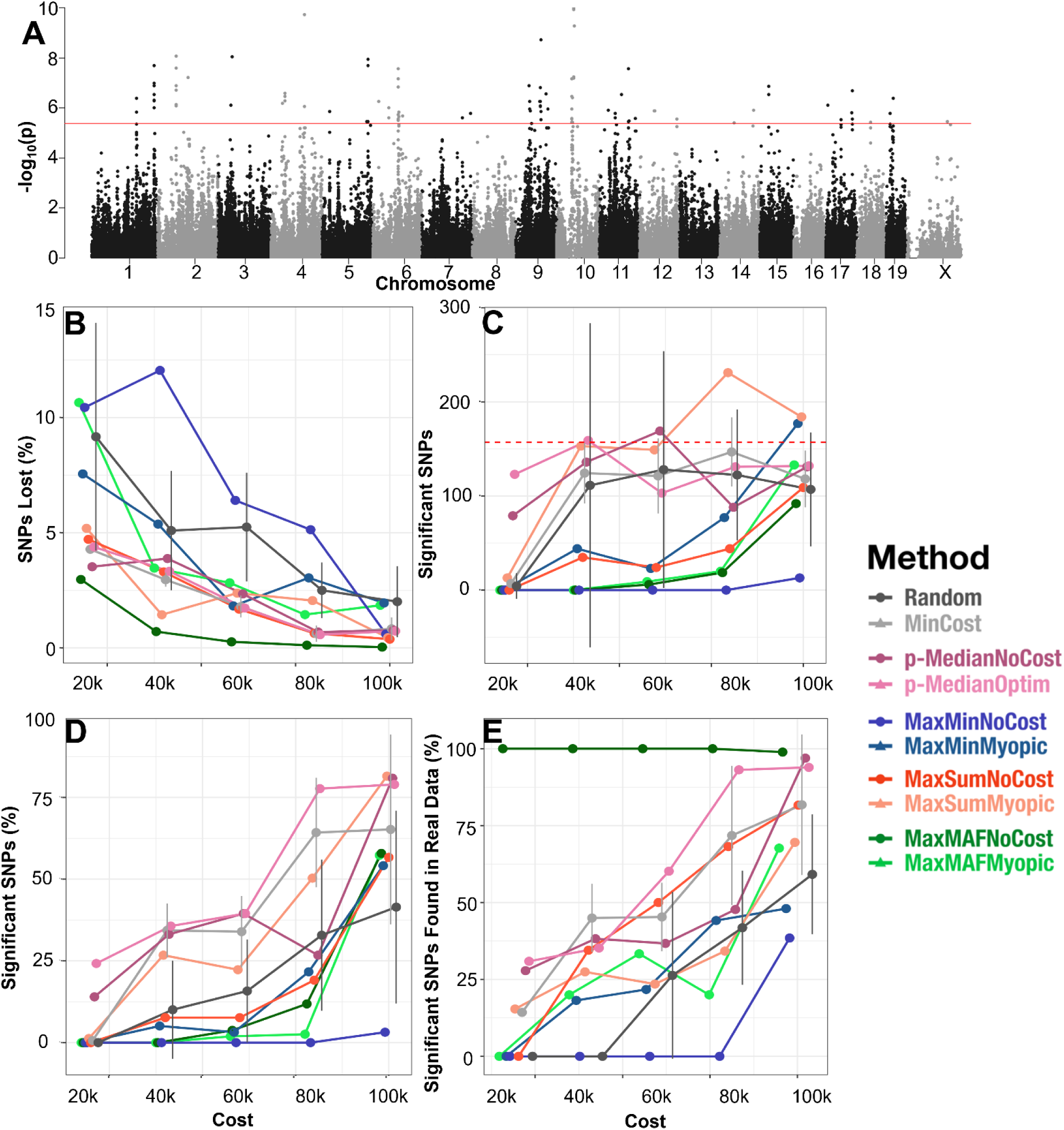
Mouse GWAS on Actual Data. Results of subsetting studies (N=10 permutations each) for real data pulled from the HMDP Heart Failure Study^12^ at $20k cost intervals from 20-100k. **A)** Manhattan plot of the results of the whole panel, red line represents the genome-wide significance threshold of a FWER of 5%^29^ **B)** Percentage of SNPs untested in the subsets because of changing MAFs. **C)** Number of significant SNPs found by each approach, the dotted red line indicates the number of significant SNPs found in the whole panel. **D)** Percentage of preserved genome-wide significant SNPs after subsetting between the subset and the full panel. **E)** The percentage of found significant SNPs which are also significant in the whole panel GWAS. Values are slightly jittered on the x axis to allow for observation of each method’s error.

In terms of total significant hits, p-MedianNoCost, MaxMinMyopic, and especially MaxSumMyopic found more significant SNPs than the original study at certain budget thresholds (Figure 6C). MaxMinNoCost, predictably, did the least well, followed by the maxMAF models, which, as before, returned the fewest number of strains (Figure S1D). Despite finding more significant SNPs than the full panel, however, MaxSumMyopic did *not* recover a majority of the full panel’s signals until the largest budgets, with p-MedianOptim and minCost leading this metric at every threshold (Figure 6D). This divergence — that is, finding many new significant SNPs that did not overlap with those found in the full dataset — hints at the possibility of inflated false-positive rates when MaxSumMyopic is used. Alternatively, selection via MaxSumMyopic may introduce a bias in population structure, causing it to no longer reflect the kinship relationships in the full panel.

Finally, MaxMAFNoCost returned very few significant hits compared to the other approaches, but with *very* high overlap with the full panel’s results: 100% of its hits were found in the original study up until the largest budget, where a single novel SNP crept in. Its marginal performance was initially unexpected. However, a more careful analysis of the significant SNPs suggested an explanation: many had MAFs of 20% or more, a regime where our simulations showed weaker performance for MaxMAFNoCost. Additionally, many SNPs had even smaller effect sizes than we had simulated, which when spread over relatively high-frequency alleles, likely further limited the efficacy of this approach.

### Future directions

We primarily considered costs by using a version of Weizman’s “myopic rule,” a greedy search that penalizes a strain’s genetic distinctiveness by its cost. Greedy searches often give reasonable solutions, but are not guaranteed to reach a global optimum for every objective. We implemented global, cost-aware optimizers using mixed-integer linear programming (MILP) for p-Median, MaxMin, and MaxSum, but only the p-Median version proved to be computationally tractable on our panel sizes. While these optimization problems are NP-hard, better heuristics and more efficient search strategies have made similar problems much more tractable^64^. It should therefore be possible to develop more efficient algorithms for balancing cost with different diversity metrics. We expect these efforts to be especially worthwhile in the case of MaxMAF, as our simulations indicate that MaxMAFMyopic’s reduced performance on mouse data can be at least partly attributed to a failure of the simple greedy search to reach a global optimum.

In the current study, we modeled the effects of selecting subsets of strains from a larger cohort based purely on cost and/or genetic diversity. Other considerations, however, can also play a role in strain selection. For example, there is a wealth of previously generated data on a number of these strains available through websites such as the Mouse Phenome Database^65^ or BacDive^49^ that may guide researchers to select certain strains with divergent phenotypes of interest as a ‘seed’ around which to grow a larger panel^65^. This approach is compatible with the selection algorithms we described in this study, which can all accommodate a specific set of starting strains, and so integrating prior knowledge about phenotypic extremes may be a productive future direction for preserving power.

Future work could focus on better understanding the tradeoffs between cost, diversity and minor allele coverage through multi-objective optimization^66^, the incorporation of kinship-aware penalties to limit potential false positives in sub-panels, and stability analyses across budgets and phenotypes to identify optimal cohorts at any given budget point.

While some power loss is unavoidable, our results do highlight that considering both cost and diversity may help mitigate this loss, allowing researchers to preserve more funding for downstream analyses or personnel costs. In particular, our results identify the cost-aware versions of MaxSum, MaxMAF, and p-Median as especially promising approaches that merit further study. At the same time, when the full panel is already diverse, our results indicate that simply maximizing the number of purchased strains can also be a reasonable strategy.

## Data and Code Availability

Mouse Diversity Array genotypes are available from the Mouse Genome Informatics website (https://www.informatics.jax.org/).

HMDP Phenotype data is available through Mendeley Resources at https://data.mendeley.com/datasets/y8tdm4s7nh/1.

All code is available at https://github.com/pbradleylab/genome_annotation_matrix.

## Acknowledgements

We wish to thank Wesley Tansey for productive conversations at the start of this project.

Funding was provided by the National Institutes of Health (R01HL162636 to CDR, R35GM151155 to PHB). CDR and PHB were also supported by startup funds from The University of North Carolina at Chapel Hill and The Ohio State University, respectively.

