## Supplemental Figures and Tables for "Optimizing strain selection for association studies under hard cost constraints"

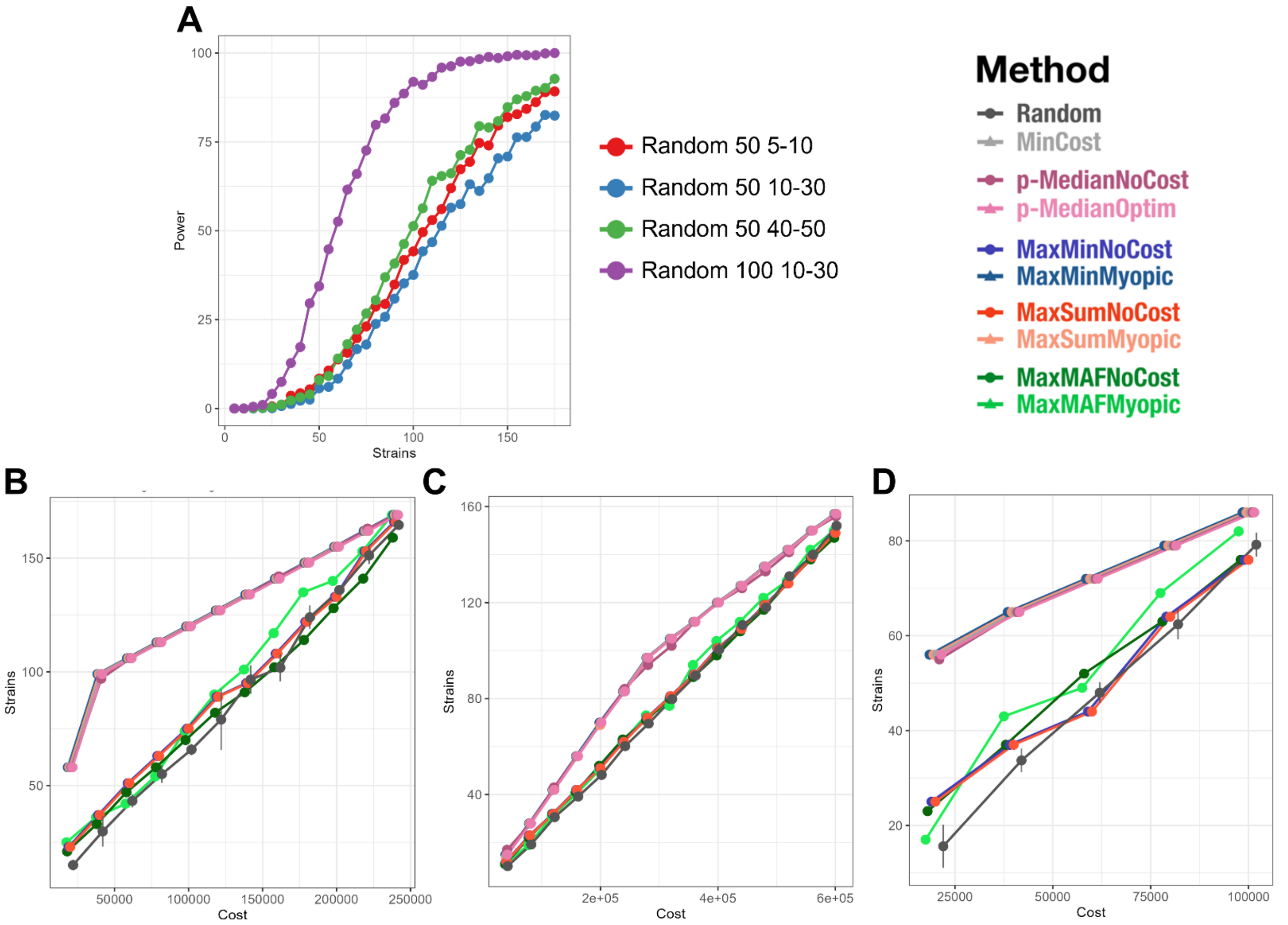


**Figure S1. Results of Random Strain Power Testing and Number of Strains Selected. A)** Results of 1,000 permutations at 5-strain intervals in the HMDP cohort for various $\chi^2$ and MAF thresholds **B)** Total number of strains selected when no additional costs are added per strain for each approach. **C)** Total number of strains selected when $2,500 is added to the cost of each strain to account for research costs **D)** Total number of strains selected for actual GWAS analysis. In all cases, x-values are shifted slightly to distinguish points, but budget cutoffs are the same as shown in Figures 4,5, and 6, respectively.

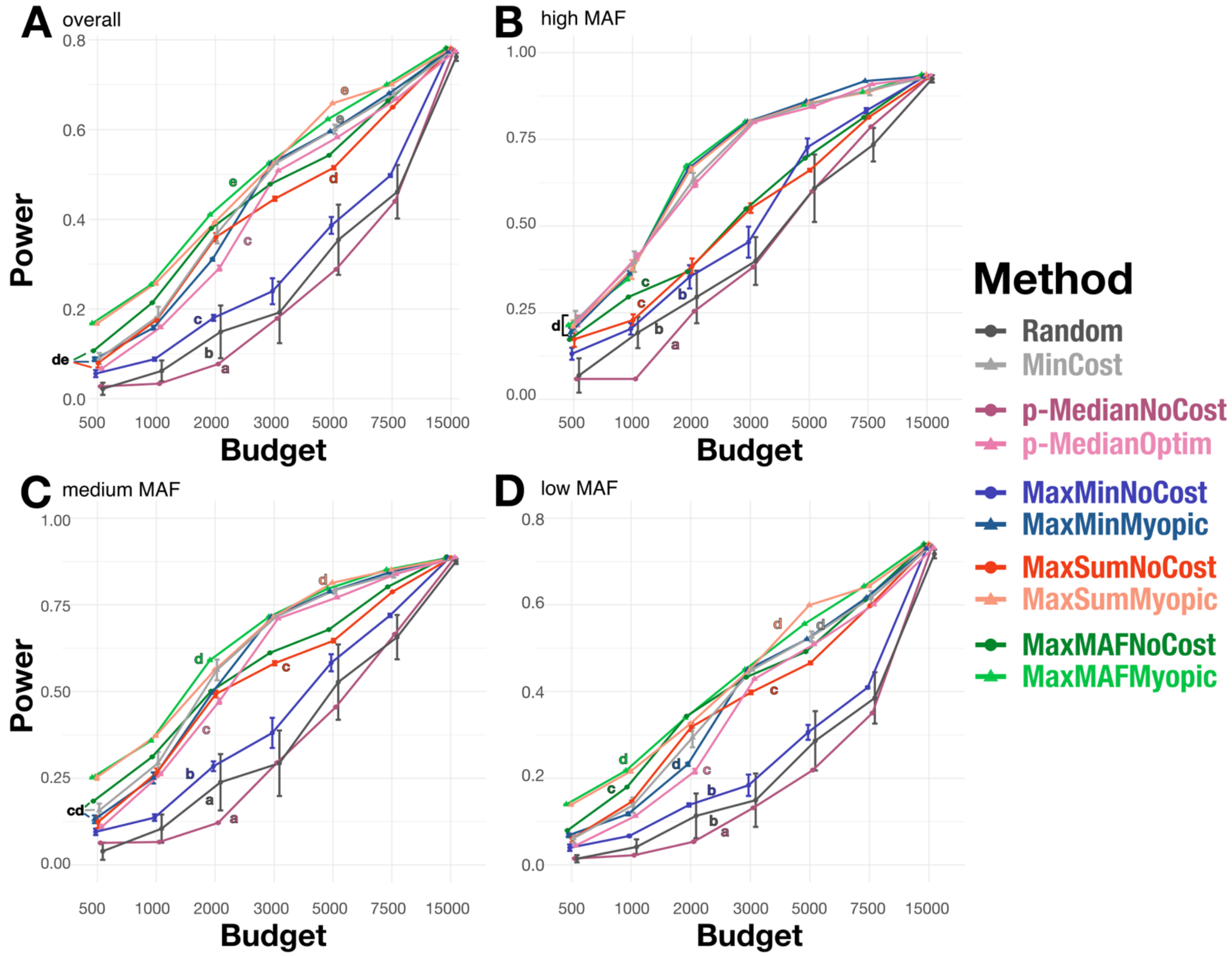


**Figure S2**. As in Figure 2 (reproduced in (A), but stratified by MAF of the causal allele. MAF ranges are $0.3 \leq \text{MAF} \leq 0.5$ (B), $0.1 \leq \text{MAF} \leq 0.3$ (C), and less than 0.1 (D). Lowercase letters (a-f) indicate significantly different groups; methods are significantly different within a panel at p=0.05 if they do not share a letter (GAM beta regression, Tukey-adjusted post-hoc pairwise tests; see Methods).

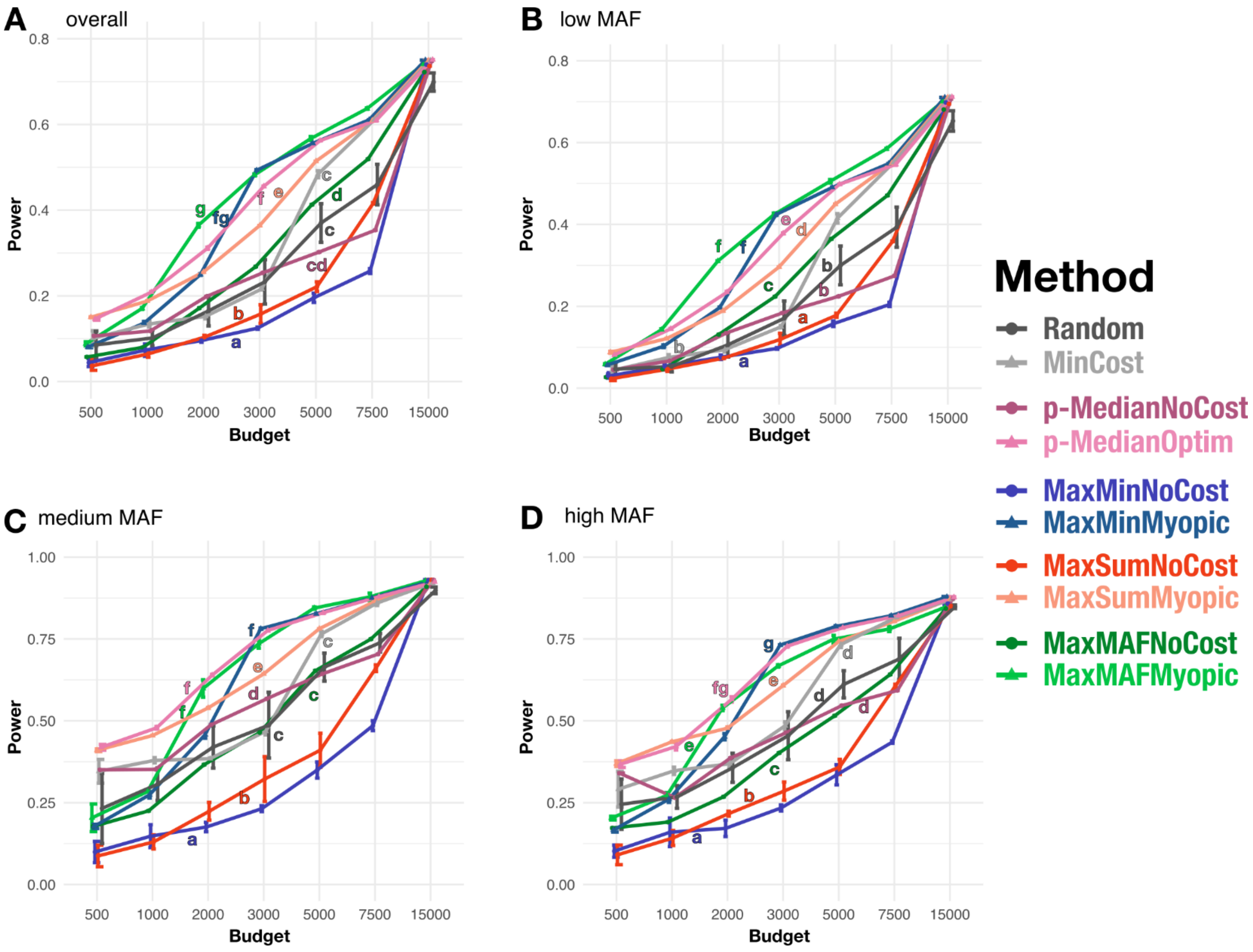


**Figure S3**. As in Figure 3 (reproduced in (A)), but stratified by MAF of the causal allele. MAF ranges are 0.3 ≤ MAF ≤ 0.5 (B), 0.1 ≤ MAF ≤ 0.3 (C), and less than 0.1 (D). Lowercase letters (a-f) indicate significantly different groups; methods are significantly different within a panel at p=0.05 if they do not share a letter (GAM beta regression, Tukey-adjusted post-hoc pairwise tests; see Methods).

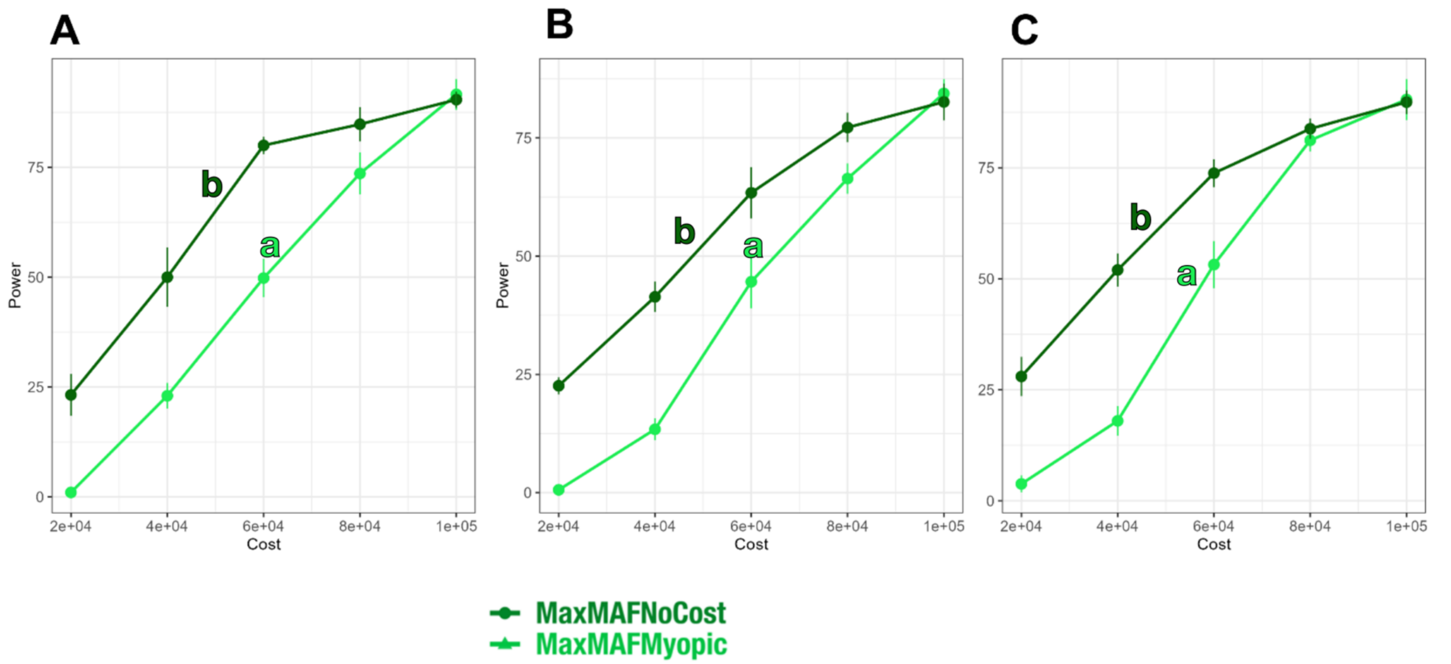


**Figure S4.** Power Comparison between maxMAFNoCost and MaxMAFMyopic when costs per strain were randomly selected from a uniform distribution ranging from \$60-\$1000. Curves marked with "a" and "b" were significantly different in a post-hoc test of a beta-regression GAM at p=0.05 (see Methods). **A)** MAF 5-10%, **B)** MAF 10-30%, **C)** MAF 40-50%.

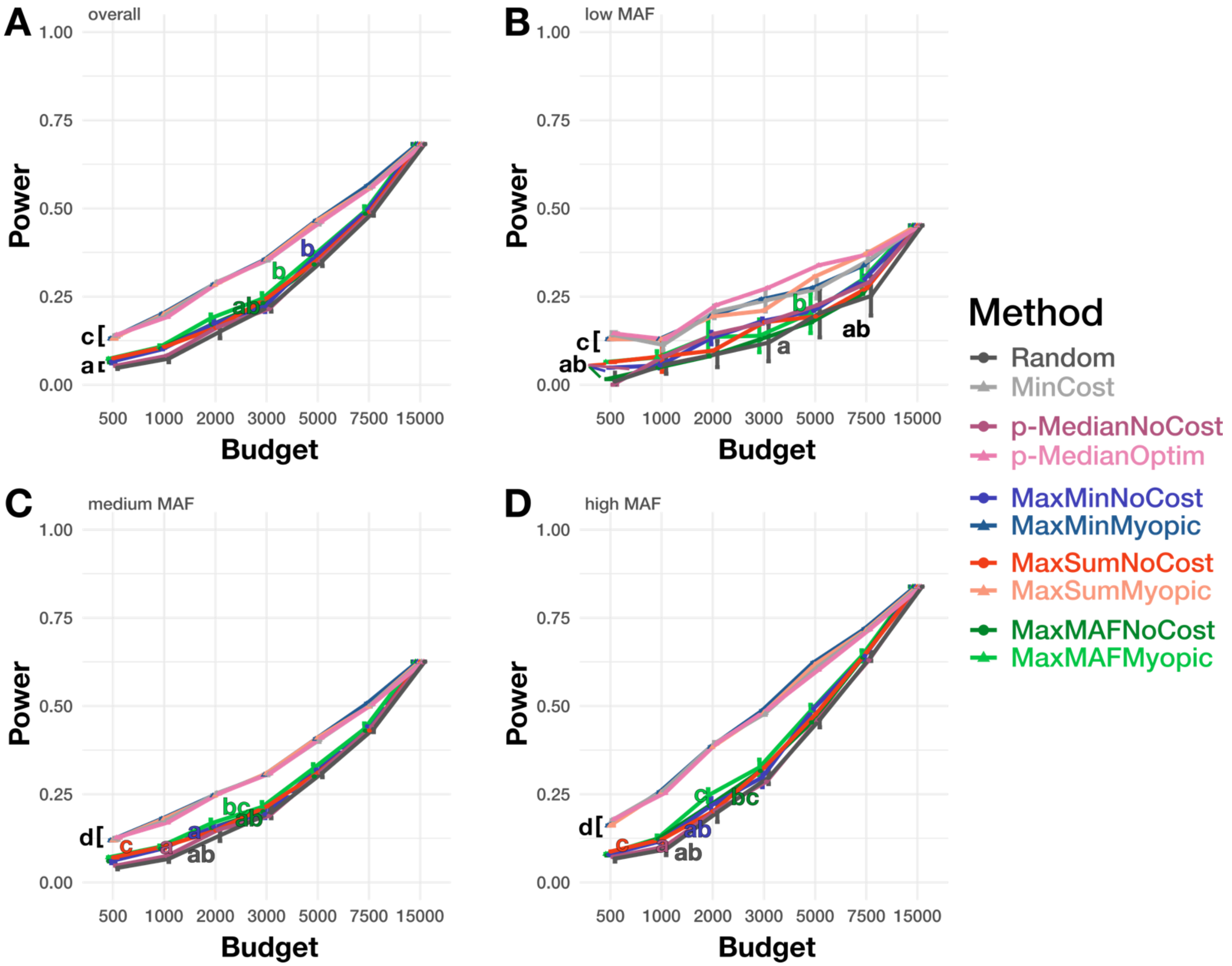


**Figure S5**. Power across budgets when "recombinant" bacterial strains were simulated from a set of six "parental" isolates. MAF ranges are $0.3 \leq$ MAF $\leq 0.5$ (B), $0.1 \leq$ MAF $\leq 0.3$ (C), and less than 0.1 (D). Lowercase letters (a-c) indicate significantly different groups; methods are significantly different within a panel at p=0.05 if they do not share a letter (GAM beta regression, Tukey-adjusted post-hoc pairwise tests; see Methods).
